# A global atlas of soil viruses reveals unexplored biodiversity and potential biogeochemical impacts

**DOI:** 10.1101/2023.11.02.565391

**Authors:** Emily B. Graham, Antonio Pedro Camargo, Ruonan Wu, Russell Y. Neches, Matt Nolan, David Paez-Espino, Nikos C. Kyrpides, Janet K. Jansson, Jason E. McDermott, Kirsten S. Hofmockel, the Soil Virosphere Consortium

## Abstract

Historically neglected by microbial ecologists, soil viruses are now thought to be critical to global biogeochemical cycles. However, our understanding of their global distribution, activities, and interactions with the soil microbiome remains limited. Here, we present the Global Soil Virus (GSV) Atlas, a comprehensive dataset compiled from 2,953 previously sequenced soil metagenomes and comprised of 616,935 uncultivated viral genomes (UViGs) and 38,508 unique viral operational taxonomic units (vOTUs). Rarefaction curves from the GSV Atlas indicate that most soil viral diversity remains unexplored, further underscored by high spatial turnover and low rates of shared vOTUs across samples. By examining genes associated with biogeochemical functions, we also demonstrate the viral potential to impact soil carbon and nutrient cycling. This study represents an extensive characterization of soil viral diversity and provides a foundation for developing testable hypotheses regarding the role of the virosphere in the soil microbiome and global biogeochemistry.

---

Viral contributions to soil ecology are largely unknown due to the extreme diversity of the soil virosphere. Despite variation in estimates of soil viral abundances (10^7^ to 10^10^ viruses per gram of soil), it is clear that soils are among the largest viral reservoirs on Earth^1–3^. Early metagenomics investigations have revealed high genetic diversity in soil viruses, with putative impacts on global biogeochemistry^1,2,4,5^. Still, less than 1% of publicly available viral metagenomic sequences are from soil^6^, reflecting the lack of knowledge about soil viruses and their ecological roles^4,7^.

High soil viral diversity may be due to the structural and/or physico-chemical heterogeneity of soil compared to other ecosystems^1,8–10^, as well as the high diversity of their microbial hosts. Indeed, soil viral abundance and composition vary with factors such as pH, temperature, moisture, chemistry, and habitat^10–12^. Much of this viral diversity is contained within DNA viruses, though RNA viruses also have the potential to influence soil processes^13,14^. While less is known about soil viral activity, a recent study of peatlands reported that close to 60% of soil viral genomes may be involved in active infections^15^, consistent with high activity observed in marine and other systems^4,16–18^.

Whether common macroecological patterns apply to the soil virosphere remains an open question; initial studies of the soil virosphere indicate that the ecology of viruses is at least partially decoupled from other microorganisms^8,10,19^. A major finding is that soil viral community turnover may occur over shorter spatial and temporal scales than microbial communities^8,10,19^. For instance, spatial viral turnover has been shown to be over 5 times higher than microbial community turnover across an 18m soil transect^8^, and only 4% of peatland ‘viral operational taxonomic units’ (vOTUs) are shared across continents^20^. Other studies note the possibility for long-distance soil viral dispersal through atmospheric^21^ or aquatic transport^22^ consistent with low turnover. These contrasting results indicate a lack of consensus surrounding the spatial and temporal patterns of soil viruses and the need for large-scale surveys of the soil virosphere.

Importantly, soil viruses can influence biogeochemical cycling, antibiotic resistance, and other critical soil functions by releasing carbon and nutrients during host infection and/or by altering host metabolism via auxiliary metabolic genes (AMGs)^9,15,18,23–28^. While soil AMG characterization is nascent^14^, marine systems demonstrate the breadth of functions ripe for discovery in soil^24^. More than 200 viral AMGs encoding functions related to carbon and nutrient cycling; stress tolerance; toxin resistance; and other processes have been detected in marine systems^24^. In contrast, only a handful of these functions have been identified as soil viral AMGs^12,14,15,22,29,30^. AMGs encoding carbohydrate metabolism in particular may be present in soils, including a few that have been experimentally validated^9,10,15,29–31^.

Accordingly, understanding the role of viruses in soil ecosystems is one of the most pressing current challenges in microbial ecology^32^. Despite the expansion of studies characterizing soil viruses^4,12,29,30^, a comprehensive description of the global soil virosphere has yet to be performed. Such a description is necessary to begin to address questions regarding the spatiotemporal dynamics, physicochemical interactions, host organisms, and food web implications of the soil virosphere. We present a meticulous compilation of the Global Soil Virus (GSV) Atlas based on previous metagenomic investigations of worldwide soils. This atlas represents the most extensive collection of soil metagenomes to date, encompassing contributions from prominent repositories, ecological networks, and individual collaborators.

## RESULTS

### Global Soil Virus (GSV) Atlas

For a description of the files contained by the GSV Atlas, please see ‘Data Availability’. We amassed 1.25 × 10^12^ of assembled base pairs (bp) across 2,953 soil samples, including 1,552 samples that were not previously available in the United States Department of Energy’s (DOE), Joint Genome Institute’s (JGI) IMG/M database (Fig. 1 and 2). These samples were screened for viruses, yielding 616,935 uncultivated virus genomes (UViGs) of which 49,649 were of sufficiently high quality for further investigation (see methods). To quantify the extent of new viral diversity encompassed by the GSV Atlas, we compared sequences from samples not already in IMG/VR to those that were previously deposited. Newly contributed sequences clustered into 3,613 vOTUs of which only 317 clustered with existing viral sequences in IMG/VR. The vast majority associations with IMG/VR were with sequences previously uncovered from soil habitats (Fig. 2b).

**Fig. 1.**
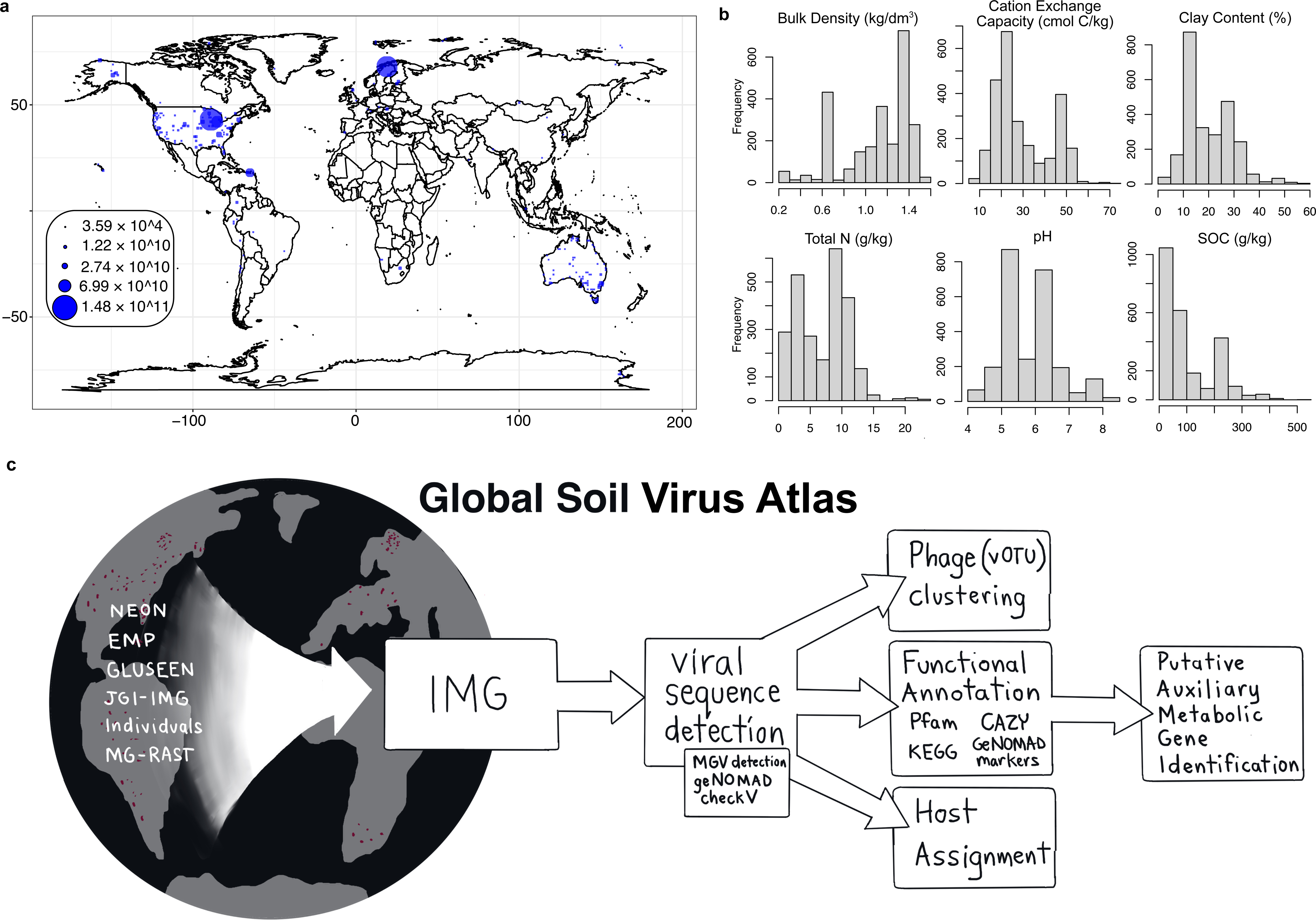
Data collection and workflow. **a**, Global distribution of samples, scaled by assembled base pairs. In order horizontally, histograms of (**b**) mean soil bulk density (kg/dm^3^), cation exchange capacity (cmol(c)/kg), clay content (%), total nitrogen content (g/kg), pH, and soil organic carbon (g/kg) associated with our samples from the SoilGrids250 database (0-5 cm). **c,** Sequence processing pipeline.

**Fig. 2.**
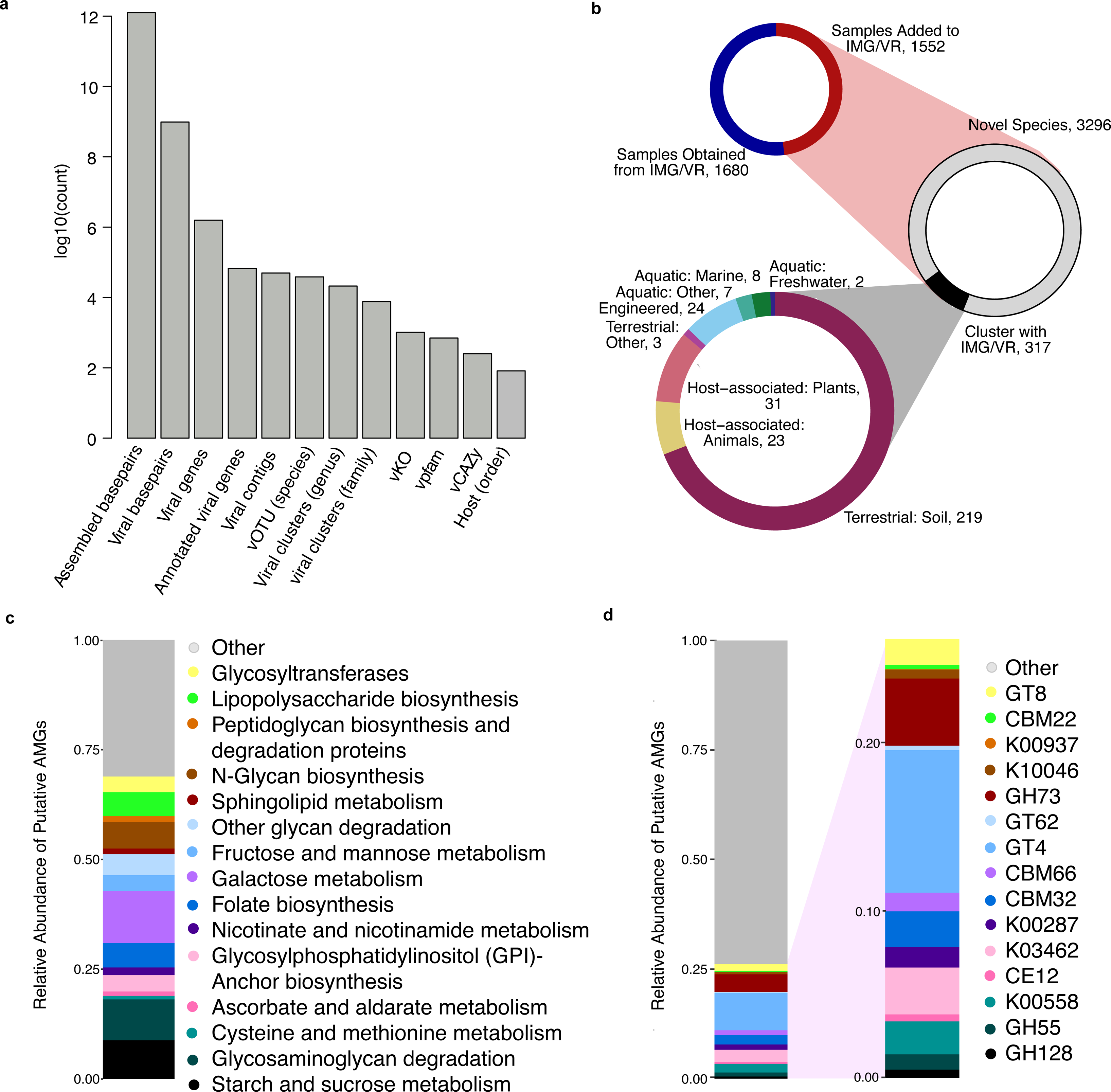
Data description. **a**, shows the count of each category across the full dataset. **b**, shows the proportion of samples obtained from IMG/VR versus the number of new samples contributed (upper left). Within new samples, we identified 3,613 vOTUs of which 317 clustered with sequences already in IMG/VR (middle). Sequences in IMG/VR that clustered with vOTUs containing new sequences were mostly associated with soil habitats (lower left). **c-d,** relative abundance of putative AMGs grouped by KEGG pathway **(c)** and by annotation **(d)**.

We also collected associated environmental parameters describing each sample from the SoilGrids250m database^33^. We assayed a wide variety of soils that ranged from bulk density of 0.24-1.56 (kg/dm^3^), cation exchange capacity of 6.8-71 (CEC; cmol(c)/kg), nitrogen content of 0.19-22.4 (N; g/kg), pH of 4.3-8.5, soil organic C of 1.9-510.9 (SOC; g/kg), and clay content of 2.7-57.1 (%) (Fig. 1).

The 49,649 UViGs of sufficient quality for downstream analysis clustered into 38,508 vOTUs at the species-like level^34^, of which 3,296 were previously unrepresented in IMG/VR (Fig. 2a-b). Only 13.9% of the GSV Atlas’ vOTUs appeared in more than one sample, and less than 1% were present in more than 5 samples. At higher taxonomic levels, we found 21,160 and 7,598 clusters at the genus and family level respectively^35^. This equates to an average of 40.01 (range: 1 - 2,124), 35.48 (range: 1 - 1,651), and 24.91 (range: 1 - 896) unique viral clusters per sample at the species, genus, and family levels. 38,278 out of 38,508 vOTUs (99.4%) had at least one member assigned to a taxon by geNomad.

We identified 1,432,147 viral genes, of which only 260,258 (∼18%) were found in at least one annotation database (1,022, 3,634, and 145 unique KO, Pfam, and CAZy annotations, Fig. 2). After filtering to putative AMGs (see methods)^30^, we found 5,043 genes that mapped to 83 KEGG pathways (1,941 KO and 3,357 CAZyme, some genes were associated with both KO and CAZyme). The median per sample putative AMG abundance (gene copies per sample) was ∼19 genes (median = 4, reflecting skewing from a few samples with high AMG abundance).

Some KEGG pathways represented by the most putative AMGs were associated with major soil carbon cycling processes (Galactose metabolism and Starch and sucrose metabolism). Likewise, at the level of gene annotations, the most common putative AMGs suggested a role for viruses in soil carbon cycles; including Glycosyltransferases 4 (GT4), glycosylhydrolases 73 (GH73), and Carbohydrate Binding Module 50 (CBM50). Other abundant KEGG pathways and gene annotations (Fig. 2c-d) included Glycosaminoglycan degradation, N-Glycan biosynthesis, Folate biosynthesis, 6-pyruvoyltetrahydropterin/6-carboxytetrahydropterin synthase (K01737), and 7-carboxy-7-deazaguanine synthase (K10026).

In contrast to the saturation observed in rarefaction curves for microbial taxonomy and genes annotated by Pfam, rarefaction curves of soil viruses for individual samples (vOTUs and and viral clusters) and their genes (annotated by Pfams) did not reach an asymptote (Fig. S1). The number of new and unique vOTUs and viral clusters at the family level (Fig. S1b and S1c) was linearly related to sequencing depth, while viral Pfams displayed slight curvature. These linear relationships were observed when considering 2TB of metagenomic sequencing––4-fold more sequencing depth than any other soil metagenome in this study and 40-fold more than the Joint Genome Institute (JGI) recommended sequencing depth for soil samples (45 GB). When considering cumulative unique attributes versus sequencing depth (Fig. S2), relationships in vOTUs and viral clusters displayed slight curvature, while viral Pfams neared saturation.

### Microbial Hosts of Soil Viruses

We connected 1,450 viruses to putative hosts of 82 different bacterial and archaeal orders with CRISPR spacers. This equates to 2.78% of QA/QC-ed viral contigs that were associated with CRISPR spacer hits; roughly 70% more host assignments than in another recent assessment^4,36^. While we observed a maximum of 73 vOTUs per host (i.e., CRISPR spacer), the mean overall vOTU per host ratio was 0.42 (median = 0), reflecting the predominance of unique host associations for individual vOTUs.

Out of 1,223 samples with at least one vOTUs assigned to a host, only 72 samples had an average of more than one host sequence per vOTU, underscoring the low abundance of detected hosts across all soils. An average of 0.64 unique host orders were detected per sample, with a maximum ratio of CRISPR spacer hits to viral sequences of 73. Further, samples with a high ratio of vOTU:host almost exclusively were matched to host sequences from a single microbial order, reflecting high phylogenetic conservation of host associations. Of the 10 samples with the highest CRISPR spacer sequence to viral sequence ratio, only 1 contained a CRISPR spacer matching more than one microbial order.

The most prevalent host taxa were distributed across distantly-related phyla, including members of prominent soil orders such as Pseudomonadales, Burkholderiales, Acidobacteriales, and Bacteroidales (Fig. 3b). The frequency of CRISPR hits associated with Acidobacteriales, Oscillospirales, Pedosphaerales, and Geobacterales were positively correlated to soil nitrogen, organic carbon, and CEC; while Enterobacterales, Obscuribacterales, Mycobacteriales, Pseudomonadales, and Streptomycetales were positively correlated to soil bulk density, and to a lesser extent, pH and clay.

**Fig. 3.**
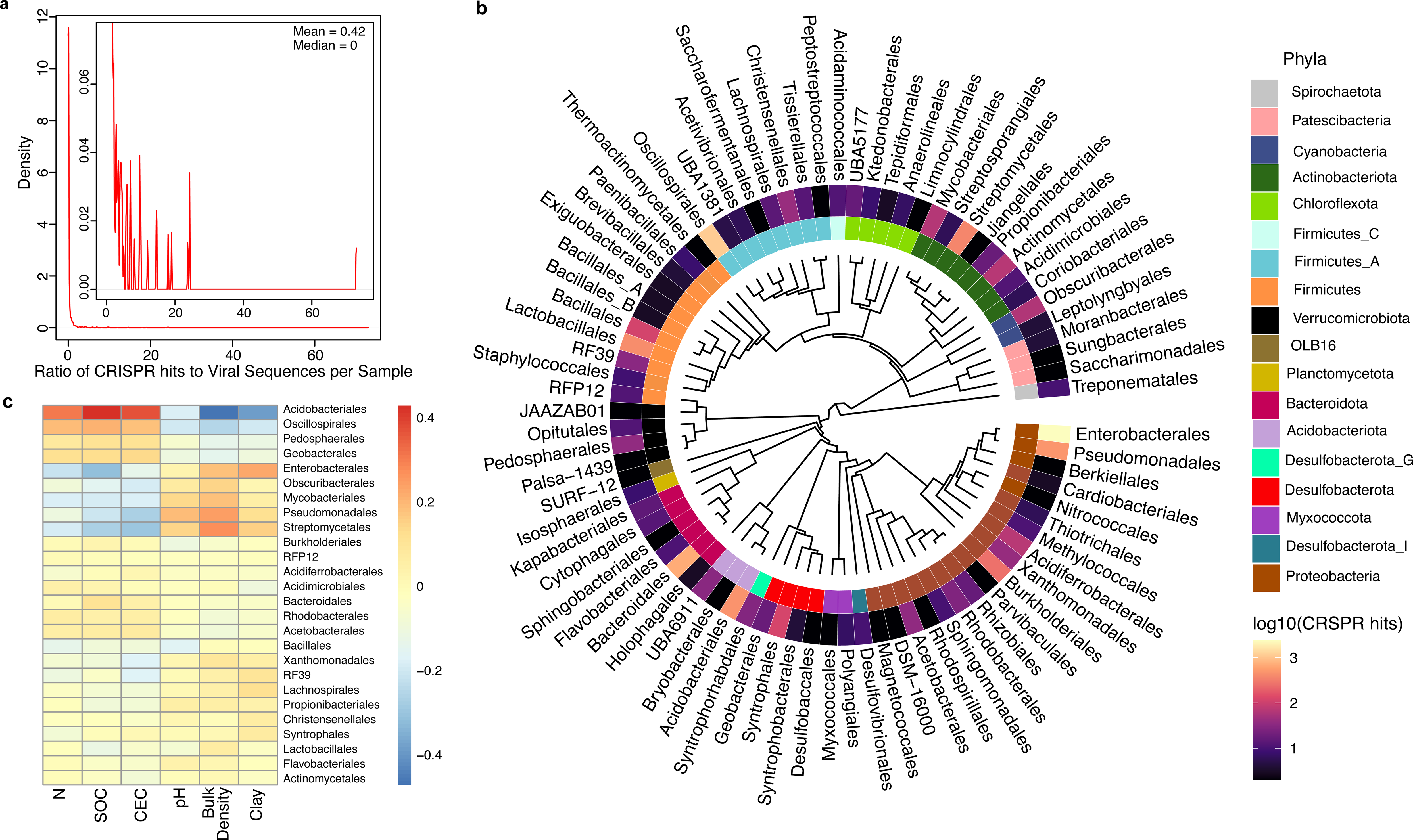
Relationships between soil viruses and their hosts. **a,** Cumulative Distribution Function plot showing the ratio of CRISPR spacer hits to viral sequences per sample. Inset shows a zoomed portion of the plot from 0 to 0.08 along the y-axis. **b,** Phylogenetic tree of bacterial hosts at the order level. Phylum taxonomy is shown in the inner circle and the abundance of CRISPR spacer hits to each order is shown in the outer circle. Two archaeal orders (Nitrososphaerales and Halobacteriales) of hosts are not shown. **c**, Correlation between frequency of CRISPR hits (defined as total CRISPR spacer hits per microbial order) and environmental parameters from SoilGrids250m. Color denotes Spearman rho. For inclusion in the heatmap, host order must be present in more than 5 samples.

### Metabolic potential encoded by the soil virosphere

Because viral gene annotations were sparsely distributed across many functions, we screened all viral genes (regardless of assignment by the AMG pipeline) against KEGG pathways to understand relationships among genes in the context of known metabolic processes. Across the entire soil virosphere, we uncovered portions of KEGG pathways that were mostly complete, including functions involved in amino acid and sugar metabolism and biosynthesis, antibacterial mechanisms, nucleotide synthesis, and other viral functions (e.g., infection strategies, Extended Data and Fig. 4). For instance, genes associated with DNA mismatch repair (map03430), homologous recombination (map03440), and base excision repair (map03410) were prevalent (Fig. S3). Folate biosynthesis was also common in the soil virosphere (map00670, map00790, Fig. 4). Bacterial secretion systems (map03070, Fig. S4), which may be evolutionarily derived from phages^37^, and the *Caulobacter* cell cycle (map04112, Fig. S5), which has a distinct division pattern^38^, were rife across soils. The GSV Atlas also contained many viral amino acid biosynthesis/degradation pathways that could be critical in viral life cycles (e.g., map00250, map00260, map00270, map00330, map00340, Fig. S6). Finally, we found nearly complete portions of energy-generating pathways including the pentose phosphate pathways and F-type ATPase-mediated portions of photosynthesis. Lipopolysaccharide pathway-related genes (LPS) that may be important as host receptors for bacteriophage and prevention of superinfection were also prevelant^39,40^.

**Fig. 4.**
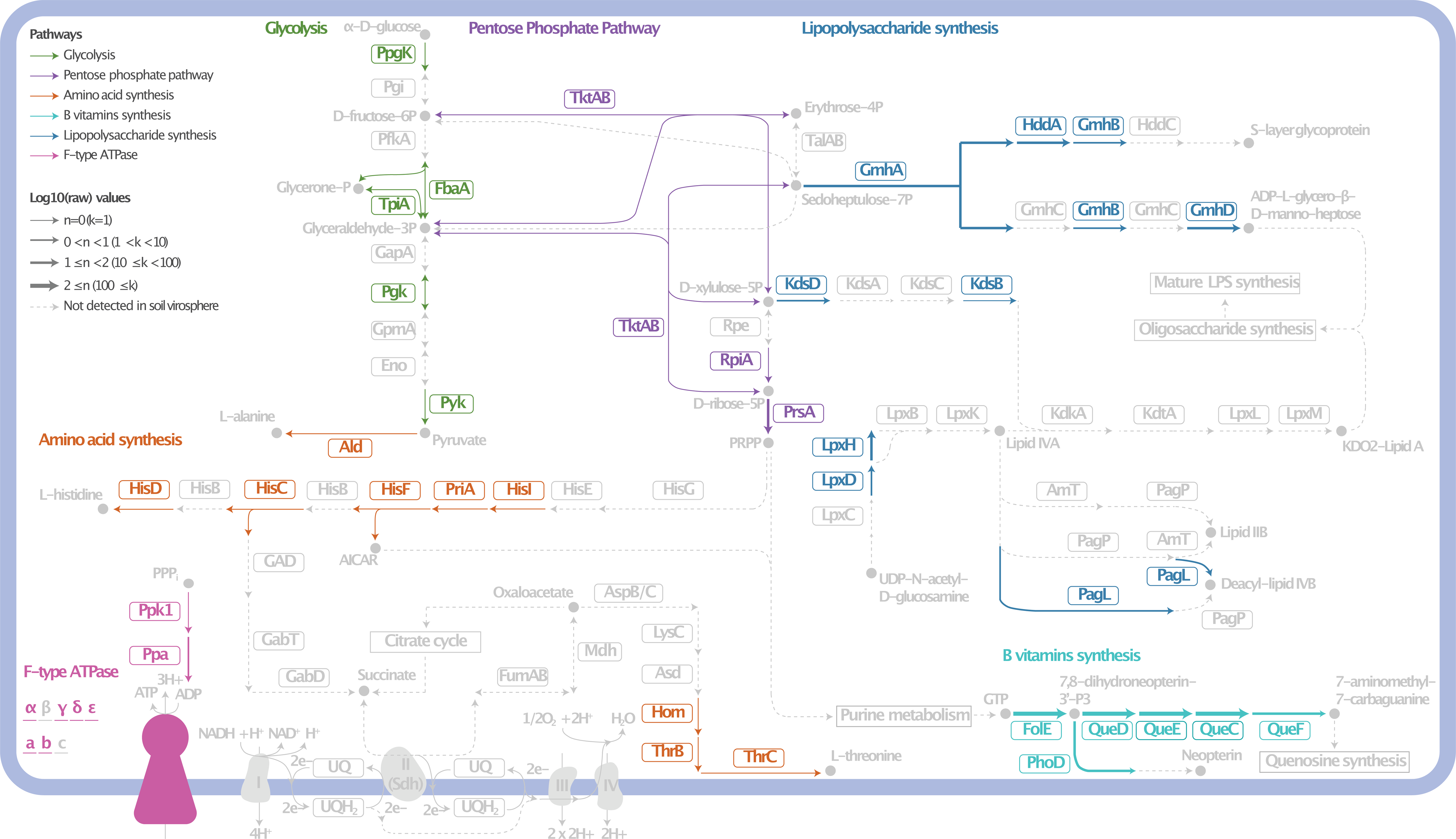
Metabolic potential encoded by the soil virosphere. A cellular diagram depicting portions of the F-type ATPase (map00190), lipopolysaccharide pathway (map00540), pentose phosphate pathway (map00030), and vitamin B- and amino acid-related KEGG pathways in the soil virosphere. Genes detected in the soil virosphere are colored according to pathway type. Undetected genes and associated metabolites (unmeasured) are grayed out. Arrow width denotes gene abundance.

## DISCUSSION

The GSV Atlas demonstrates the immense, unexplored taxonomic and functional diversity of the soil virosphere. Viral diversity in the GSV Atlas appeared to be largely distinct from other global habitats. Nearly 80% of GVS Atlas sequences that clustered with existing sequences in IMG/VR were attributed to soil or soil-like habitats [i.e. ‘other terrestrial’ or ‘plant-associated’ (rhizosphere)], underscoring the unique composition of the soil virosphere relative to more well-studied marine and human environments. Additionally, few shared vOTUs and viral clusters between samples may indicate high spatial turnover (i.e., changes in soil virosphere composition through space). Recent studies have estimated soil viral diversity is high, both relative to other viral habitats and relative to soil microbial diversity^7,8,10,22^. However, these estimates have been limited by copious viral and microbial ‘dark matter’ for which no functional or taxonomic assignment is known^14,23,32^. Towards this end, the diversity encompassed by the GSV Atlas can serve as a community resource for characterizing this unknown fraction of the soil virosphere.

Analysis of the GSV Atlas suggests that extreme spatial heterogeneity may be a key feature of the soil virosphere at the global scale. While rapid viral spatial turnover was recently observed across short spatial scales (<10-20m)^8,10^, there has been no such demonstration of viral biogeography across the world. We propose that high rates of spatial turnover could result from low dispersal rates or distinct temporal dynamics of viral communities relative to other organisms. For example, while dormant microorganisms and relic DNA can persist for months or more^41–43^, the burst of viral cells associated with active infections may generate short-lived pulses of distinct viral communities that do not contribute to relic DNA due to their comparatively small genome sizes versus microorganisms. Additionally, the apparent discrepancies between microbial and viral dispersal processes could be due to the presence of free viruses that are not actively involved in microbial infection^14^, smaller viral genomes that could facilitate physical protection, differences in traits that facilitate dispersal between viruses and microorganisms, variation in bioinformatic pipelines, and/or other ecological differences between viruses and microorganisms.

Together, these factors make characterizing the soil virosphere a challenge for the coming decade. When examining individual soil samples, the number of new and unique viral attributes (e.g., vOTUs, clusters, Pfams) was linearly related to sequencing depth, suggesting that new viral discoveries are likely to continue with increasingly deep sequencing (Fig. S1). This contrasts with rarefaction curves of the soil microbiome and of microbial hosts of soil viruses, which both asymptoted well before sequencing depths of typical soil microbial investigations. Still, when looking at the cumulative number of unique viral attributes detected in all samples collectively (Fig. S2), many viral attributes began to saturate with sequencing depth. This suggests that while individual samples do not capture soil viral diversity, we can begin to constrain the extent of diversity when sequences from thousands of existing samples are aggregated.

Functional diversity encoded by the GSV Atlas revealed the potential for soil viruses to impact biogeochemical cycles; in particular by supporting organic matter decomposition. KEGG pathways and gene annotations represented by the most putative AMGs were related to the metabolism and/or production of sugars common to soils including sucrose, mannose, glucosamine, and maltose^44,45^; as well as the decomposition of chitin – one of the most abundant carbon molecules in soil^46^. Our results are consistent with previous work from single locations that have hinted at a wide range of possible soil viral AMGs, including glycoside hydrolases, carbohydrate esterases, and carbohydrate-binding modules^15,23,31^. Given that a large proportion of soil microorganisms are infected by viruses at any given time^47^, AMGs encoded by soil viruses have the potential to impact global biogeochemical cycles^15,22,23,31^. The thousands of putative AMGs identified here represent the most extensive survey to date and further impress the importance of the soil virosphere as a reservoir for biogeochemical potential.

Unraveling relationships between viruses and their host communities is imperative to understanding the impact of the virosphere on soil processes. Host presence should be tightly coupled to viral abundance, and in turn, these linkages are mediated by spatial, temporal, and environmental factors^15,48,49^. These linkages are also dependent on viral host range (i.e., host specificity) –– higher host specificity should lead to stronger coupling between microbial and viral abundance and community composition. Viral host specificity is also associated with ecological factors that impact microbial community composition and may result in tradeoffs between viral growth and the breadth of the host range^11,50–52^. Across the GSV Atlas, there were few hosts per vOTU on average (mean = 0.42), and of vOTUs associated with multiple host sequences, the vast majority were linked to multiple hosts of the same phylogenetic clade. While high host specificity has historically been the prevailing paradigm, our work contrasts recent studies suggesting that some soil viruses may have broader host ranges than viruses in other habitats^53,54^.

The ultimate impact of viral predation on soil functions is at least partially associated with the taxonomic distribution of hosts and their variation across soil habitats. Host sequences spanned nearly every major soil microbial clade, consistent with other recent studies (Fig. 3)^22,23,31^. This taxonomic breadth suggests a role for the soil virosphere in most soil habitats. Moreover, some hosts were susceptible to changes in the environment, which may reflect environmental filtering on host communities (which in turn, determine the amount and type of viruses present) or on viruses directly which subsequently impacts host community composition^15,27,55,56^. Viral infections have been previously linked to soil parameters including moisture^12,30^ and carbon and nitrogen content^9^. In our analysis, bulk density may serve as a proxy for hydrologic connectivity in the soil matrix. For example, low hydrologic connectivity may create ‘spatial refuges’ for soil bacteria from viral infections^8^, influence the virus-host encounter rates, and thus structure the soil virosphere and its hosts. Nutrient amendments are also considered to be drivers of the soil virosphere, supporting the relationship we observed between carbon, nitrogen, and host taxa.

When examining the functional potential of the soil virosphere, we detected many hallmarks of viral activity––including genes associated with cell lysis, DNA repair/replication, and other infection signatures––and viral amino acid biosynthesis/degradation pathways that could be critical in viral life cycles (Fig. 4, Fig. S3-S6). The prevalence of viral genes associated with central microbial functions highlights the potential importance of viral activity in soils and the need for targeted approaches to quantify the extent and impact of viral gene expression. For instance, folate and other B vitamins may be logical targets for pathogens as they are key to bacterial growth(map00670, map00790, Fig. 4)^57,58^. Type IV secretion systems can be used by bacteria to secrete toxins^59^ or as a method for DNA transfer through membranes.^60^ The *Caulobacter* cell cycle (map04112, Fig. S5) is another promising indicator of viral infections due to its distinct cell division process^38^. Finally, amino acids are building blocks for cellular material and also support soil biogeochemical cycles, as they can enhance carbon cycling through priming effects and/or enhanced nutrient availability^61,62^ (e.g., map00500, map00052, map00051, Fig. 4). Collectively, these pathways demonstrate several possibilities for soil viral impacts on processes that are central to microbial metabolism and biogeochemical cycling of elements in soil.

Beyond these pathways, we highlight three KEGG pathways with near-complete portions represented in the GVS Atlas: F-type ATPase (map00190), pentose phosphate pathway (map00030), and LPS (map00540). Five of seven subcomponents of the F-type ATPase were detected in the soil virosphere, while no V- or A-type ATPases were found. Given the evolutionary similarities between V- and F-type ATPase in particular^63^, the lack of any V- or A-type ATPase components is notable in light of the near complete F-type ATPase. Though there is some basis for F-type ATPases in viral replication^64^, we also note the possible involvement of F-type ATPase in photosynthetic energy generation^65^. Given the prominence of photosynthetic marine AMGs^26,66^, we highlight the possibility of a viral F-type ATPase as a soil AMG. The pentose phosphate pathway is also a prevalent and important AMG found in marine ecosystems, where viral infection diverts carbon towards the pentose phosphate pathway as an ‘express route’ of energy generation, at the expense of host carbon metabolism (reviewed in ^66^). Finally, we observed nearly complete LPS pathways in the GSV Atlas. Phages often carry depolymerases and other enzymes that target LPS or similar outer membrane components to facilitate binding and entry^39^. However, the representation of the LPS synthesis pathway by putative soil AMGs indicates that phage may work to change the function of the pathway post-infection, potentially to prevent superinfection^40^. Collectively, we propose that F-type ATPase, pentose phosphate pathway, and LPS may be interesting pathways for more targeted investigations into the role of the virosphere in soil microbiome function.

The field of soil viral ecology is poised for rapid expansion; yet several challenges remain in fully characterizing soil viral diversity and function. Overcoming these methodological and ecological hurdles will require broad participation from global researchers. Below, we present a summary of issues, from our perspective, facing the current generation of soil viral ecologists and suggestions for surmounting them.

First, we propose methodological investments to improve viral detection and resolve genomic ‘dark matter’. Metagenomic sequencing enables the detection of thousands of viruses per soil sample, yet the number of viruses detected in soil metagenomes has remained relatively flat over time^4^. In part, this is because soil metagenomics from shotgun sequencing is highly fragmented, leading to lower quality UViGs^67,68^. Identifying novel viral sequences and assigning viruses to microbial hosts are also limited by the extent of our knowledge of viral diversity; thus expansion of the known virosphere is needed. Technical advances may improve soil virus identification and host-linkage predictions from shotgun metagenomics, long read sequencing, and/or targeted sequencing approaches. Promising new methods include experimental verification of viral activity^29^, size fractionation (‘viromics’)^7,8,15^, viral isolation^69^, optimized viral nucleic acid extraction^70^, microscopy^29^, combined metagenomic assembly^4^, and long-read and/or single-cell sequencing^71,72^.

Knowledge about soil viral diversity and function is also limited by gaps in field and laboratory experiments. The GSV Atlas demonstrates that extensive, spatially-explicit sampling is needed to capture the high spatial turnover of the soil virosphere. The spatial coverage of most ‘global’ ecological studies, including this one, often suffers from large data gaps^73^. Concerted efforts are needed to sample wide spatial domains, including historically undersampled regions, given the high viral diversity uncovered by the GSV Atlas. Expansion of the known virosphere in this way will also help to facilitate tool development. Although we did not assess temporal dynamics, temporally-explicit approaches are likewise needed to characterize temporal dynamics in soil viral communities. Further, our functional annotation of viral contigs revealed diverse genes associated with functions relevant to both viral and microbial communities, and it is impossible to know the true functions of viral genes without targeted functional assays. We therefore propose that experiments targeting the expression and auxiliary metabolic function of viral genes are needed to properly assess AMGs in viral communities.

Finally, we still know relatively little about the ecological drivers of soil virus distribution or how to represent these mechanisms in process-based models. Extreme soil virosphere diversity renders some common microbial ecology statistical methods unfeasible, including those often used to test ecological principles (e.g., ordinations, distance-decay, richness, etc.). This highlights the need for innovative statistical approaches to interpret the soil virosphere and to develop new theories surrounding their ecological roles. These advances can help aid development of process-based models, which have made tremendous improvements in representing soil carbon cycles but are missing dynamics involving the soil virosphere.

The GSV Atlas is a new public resource that can help generate hypotheses and provide insight into some of the most pressing challenges in soil viral ecology. We uncovered 616,935 UViGs from global soil samples to show the extreme diversity, spatial turnover, and functional potential of the soil virosphere. This includes a wide taxonomic array of microbial hosts of soil viruses, key functions associated with soil carbon cycles, and an assortment of viral metabolisms that may be critical to deciphering viral ecological principles in the soil ecosystem. We specifically highlight F-type ATPase, the pentose phosphate pathway, and LPS-related genes, as well as enzymes involved in carbohydrate metabolism, as fruitful areas for further investigation. Our work scratches the surface of the soil virosphere and serves as a basis for tool, theory, and model development to further advance soil ecology, biogeochemistry, ecology, and evolution.

*Correspondence should be addressed to*.

## Acknowledgements

This work was supported by the U.S. Department of Energy (DOE), Office of Biological and Environmental Research (BER) as part of BER’s Genomic Sciences Program (GSP) under FWP 70880. A portion of this work was conducted by the U.S. Department of Energy Joint Genome Institute (https://ror.org/04xm1d337), a DOE Office of Science User Facility, is supported by the Office of Science of the U.S. Department of Energy operated under Contract No. DE-AC02-05CH11231. We thank Young Song for his help in data visualization.

## Author Contributions

EBG, RW, JKJ, DPE, NCK, and JEM conceived of this project. EBG, RW, APC, RYN, MN, and AC conducted data analysis. EBG, RW, APC, RYB, NCK, JKJ, KSH, and JEM contributed to manuscript drafting and revisions. The Soil Virosphere Consortium contributed metagenomic data and reviewed the manuscript.

## ONLINE METHODS

### Data Collection and Curation

We collected a total of 2,953 soil metagenomic samples from soils from major repositories and ecological networks including the JGI Integrated Microbial Genomes and Microbiomes (IMG/M) platform, MG-RAST metagenomics analysis server, Global Urban Soil Ecological Education Network (GLUSEEN), Earth Microbiome Project (EMP), and National Ecological Observatory Network (NEON) plus submissions from individual collaborators. This included 1,552 samples not previously included in IMG/M (Fig. 1 and 2). All dataset authors were contacted for data reuse permissions.

For samples collected via JGI IMG/M, we retrieved all studies with GOLD^74^ ecosystem type of “Soil” as of August 2020. We manually curated metagenomic sequences to remove misclassified data as follows. We removed samples from studies with the following: (1) GOLD ecosystem types: Rock-dwelling, Deep subsurface, Plant litter, Geologic, Oil reservoir, Volcanic, and Contaminated; (2) GOLD ecosystem subtypes: Wetlands, Aquifer, Tar, Sediment, Fracking Water, and Soil crust; (4) words in title: wetland, sediment, acid mine, cave wall surface, mine tailings, rock biofilm, beach sand, Petroleum, Stalagmite, Subsurface hydrocarbon microbial communities, Vadose zone, mud volcano, Fumarolic, enriched, Composted filter cake, Ice psychrophilic, oil sands, groundwater, Contaminated, rock biofilm, Deep mine, coal mine fire, Hydrocarbon resource environments, Marine, enrichment, groundwater, mangrove, saline desert, Hydroxyproline, Rifle, coastal, compost, biocrust, crust, Creosote, soil warming, Testing DNA extraction, and/or Agave; (5) GOLD geographic location of wetland; and (6) GOLD project type of Metagenome - Cell Enrichment. Additionally, sample names that indicated experimental manipulation (e.g., CO_2_ enrichment or nitrogen fertilization) or were located in permafrost layers were manually excluded. This resulted in 1,480 curated metagenomes from publicly available data in IMG/M.

After collating samples from JGI IMG/M and the newly collected samples from external networks and collaborators, the final dataset consisted of 2,953 soils with 2,015,688,128 contigs, representing 1.2 terabases of assembled DNA sequences.

In parallel, we retrieved mean values for soil parameters from the SoilGrids250m database from 0-5 cm^33^. SoilGrids250m is a spatial interpolation of global soil parameters using ∼150,000 soil samples and 158 remote sensing-based products. Here, we focus on six parameters often associated with soil microbial communities: bulk density, cation exchange capacity, nitrogen, pH, soil organic carbon, and clay content. Because our focus on spatial dynamics and soils were collected at various times, we did not include temporally dynamic variables such as soil moisture or temperature in our set of environmental parameters, though we acknowledge they may have profound impacts on the soil virosphere.

### Assembly and Annotation of Samples added to IMG/M

To standardize data analysis across all samples, the 1,552 soil metagenomic samples not collected from IMG/M were analyzed using the JGI’s Metagenome Workflow ^75^. In brief, samples were individually assembled using MetaSpades v3.1. 1,476 of the 1,552 assembled soil samples passed default quality control thresholds^76^, yielding 133 gigabases of assembled DNA in 241,465,924 contigs. Additionally, three very large metagenomes (>1TB each) were assembled separately due to computational limitations in standard workflows^77^. The resulting assemblies were assigned GOLD identification numbers and imported into IMG/M and processed using IMG/M Metagenome Annotation Pipeline v5.0.0 to align with data obtained directly from IMG/M^75^.

### Virus identification, clustering, and host prediction

We performed an initial identification of viral contigs using using a modified version of the IMG/VR v3’s virus identification pipeline (code available at https://github.com/snayfach/MGV/tree/master/viral_detection_pipeline)^35,36^. The pipeline identifies viruses based on the presence of 23,841 virus protein families, 16,260 protein families of microbial origin from the Pfam database^78^ and VirFinder^79^ to identify putative viral genomes in contigs that were at least 1 kb long. During the course of this study, geNomad v1.3.3^80^, a tool for virus identification with improved classification performance was released and incorporated into our pipeline to improve prediction confidence and perform taxonomic assignment. We further processed predicted viral sequences using CheckVv1.0.1 (database version 1.5)^81^ to assess the quality of the viral genomes. As this study focused on non-integrated virus genomes, contigs that were flagged by either geNomad or CheckV as proviral were discarded. From the remaining contigs, virus genomes were selected using the following rules: (1) contigs of at least 1kb with high similarity to genomes in the CheckV database (that is, that had high- or medium-quality completeness estimates) or that contained direct terminal repeats were automatically selected; (2) contigs longer than 10kb were required to have a geNomad virus score higher than 0.8 and to either encode one virus hallmark (e.g., terminase, capsid proteins, portal protein, etc.), as determined by geNomad, or to have a geNomad virus marker of at least 5.0; (3) contigs shorter than 10 kb and longer than 5 kb were required to have a geNomad virus score higher than 0.9, to encode at least one virus hallmark, and to have a virus marker enrichment higher than 2.0. This resulted in 49,649 viral contigs that we used for downstream analysis. All viral contigs are available at: https://doi.org/10.25584/2229733.

Viral genomes were clustered into viral operational taxonomic units (vOTUs) following MIUViG guidelines (95% average nucleotide identity, ANI; 85% aligned fraction^34^). In brief, we performed an all-versus-all BLAST (v2.13.0+, ‘-task megablast -evalue 1e-5 -max_target_seqs 20000’) search to estimate pairwise ANIs and AFs, as described in Nayfach et al.^81^ and employed pyLeiden (available at: https://github.com/apcamargo/pyleiden) to cluster genomes, using as input a graph where pairs of genomes that satisfied the MIUViG criteria were connected by edges. Viruses were also grouped at approximate genus-level (40% AAI — average amino acid identity; 20% shared genes) and family-level (20% AAI; 10% shared genes) clusters using DIAMOND^82^ for protein alignment and Markov Cluster Process (MCL)^83^ for clustering^35^.

Viral sequences were assigned to putative host (bacterial and archaeal) taxa through matches to a previously described database of CRISPR spacers of 1.6 million bacterial and archaeal genomes from NCBI GenBank and MAGs (release 242; 15 February 2021)^84–88^. Sequences of viral genomes were queried against the spacer database (https://portal.nersc.gov/cfs/m342/crisprDB) using blastn (v2.9.0+, parameters: ‘-max_target_seqs = 1000 -word_size = 8 -dust = no’). Only alignments with at least 25 bp, less than 2 mismatches, and that covered ≥95% of the spacer length were considered. Viral sequences were assigned to the host taxon at the lowest taxonomic rank that had at least two spacers matched and that represented >70% of all matches.

### Potential Auxiliary Metabolic Gene (AMG) Prediction

We leveraged an intermediate output of geNomad (v1.3.3)^80^ (‘genes.tsv’) to screen putative AMGs on the detected viral contigs. Proteins of the viral contigs were annotated by virus and host-specific markers implemented in geNomad. The identified viral hallmark (e.g., terminase, major capsid protein) and non-hallmark proteins were labeled as ‘VV-1’ and ‘V*-0’ in geNomad output, respectively. The rest of the viral proteins of the detected viral contigs that were annotated as non-virus-specific or unclassified were then classified into five categories of putative AMGs based on the presence of viral hallmark or non-hallmarks up- or down-stream as mentioned previously^30^. The AMGs with both virus-specific genes (‘VV-1’ or ‘V*-0’) were retained for the following analysis. To improve the functional annotations of the putative AMGs and highlight the viral potentials of metabolizing carbohydrates and glycoconjugates, the AMG proteins were also annotated by Carbohydrate-Active enZYmes (CAZY) Database and Kyoto Encyclopedia of Genes and Genomes (KEGG) database using the default settings in addition to the functional annotation databases implemented in geNomad. The putative AMG was assigned to the functional annotation with the highest bitscore (e.g,. duplicate annotations were not allowed). Following Hurwitz and U’Ren^66^ and Hurwitz et al^89^, we further screened putative AMGs to remove genes not found in KEGG pathways. Additionally, in recognition of the ambiguity in distinguishing genes encoding auxiliary metabolic functions versus core metabolic processes^66^, we discuss the resulting set of genes presented here as ‘putative AMGs’.

### Statistical Analysis

All statistical analyses and data visualizations were performed using *R* v4.1.0^90^. We used the following packages for data manipulation and visualization: ggplot2^91^, reshape2^92^, pheatmap^93^, Hmisc^94^, ggpubr^95^, RColorBrewer^96^, maps^97^, stats^98^ geosphere^99^, plyr^100^, dplyr^101^, and stringr^102^. Packages pertaining to specific analyses are listed below.

We generated rarefaction curves for individual samples and for cumulative sequencing depth (Fig. S1 and S2) using the ‘phyloseq’ package ^103^ and custom R plots, respectively. Samples containing less than five vOTUs, viral clusters, or viral Pfams; or less than 100 CRISPR-spacer-based host taxon, microbial Pfams, or microbial taxa were removed for visual clarity. Removed samples followed the same general trends as presented in Fig. S1. To visualize saturation across cumulative sequencing depth (Fig. S2), we ordered samples from lowest to highest total assembled bp and progressively added them along the x-axis. On the y-axis, we plot the associated cumulative number of unique attributes.

A phylogenetic tree of CRISPR-spaced-based host taxon was generated at the order-level using phyloT v2^104^, an online tree generator based on the Genome Taxonomy Database. Then, we visualized the tree in R using the packages ‘ggtree’^105^, ‘treeio’^106^, and ‘ggnewscale’^107^. To examine relationships between common microbial hosts of soil viruses and soil properties, we first downloaded data describing bulk density, cation exchange capacity, nitrogen, pH, soil organic carbon, and clay content from the SoilGrids250m database^33^ using the ‘soilDB’ package^108^. Mean values of soil properties from 0-5 cm were correlated to the total number of CRISPR spacer hits per microbial ordered using Spearman correlation.

Finally, we mapped genes detected across the entire soil virosphere (i.e., all samples combined) to their corresponding KEGG pathways using the ‘pathview’ package in R^109^. Gene abundances were converted by log base 10 for visualization.

## DATA AVAILABILITY

The GSV Atlas is available for download at: https://doi.org/10.25584/2229733. It includes all UViGs regardless of quality (File 1, 616,935 UViGs), a file containing data associated with each contig that passed QA/QC (File 2, 49,649 contigs), predicted viral protein sequences (File 3, 402,882 predicted protein sequences), a file containing data associated with each gene (File 4, 1,432,147 genes), geographic and physicochemical data of the curated soil samples (File 5, 2,953 samples), and a readme file (File 6).

## CODE AVAILABILITY

Code for sequence processing is described in Nayafch et al.^35^ and is available in the materials associated with those publications. Github repositories associated with this publication are available at: https://github.com/snayfach/MGV/tree/master/viral_detection_pipeline and https://github.com/apcamargo/pyleiden). All other code is available upon request.

## CONSORTIUM AUTHORS

### Name; Affiliation; E-mail; ORCID

Emily B. Graham; Biological Sciences Division, Pacific Northwest National Laboratory, Richland, WA USA; School of Biological Sciences, Washington State University, Pullman, WA; 0000-0002-4623-7076

Antonio Pedro Camargo; US Department of Energy Joint Genome Institute, Lawrence Berkeley National Laboratory, Berkeley, CA, USA; 0000-0003-3913-2484

Ruonan Wu; Biological Sciences Division, Pacific Northwest National Laboratory, Richland, WA USA; 0000-0001-9466-4462

Russell Y. Neches; US Department of Energy Joint Genome Institute, Lawrence Berkeley National Laboratory, Berkeley, CA, USA; 0000-0002-2055-8381

Matt Nolan; US Department of Energy Joint Genome Institute, Lawrence Berkeley National Laboratory, Berkeley, CA, USA; 0000-0002-9532-0853

David Paez-Espino; US Department of Energy Joint Genome Institute, Lawrence Berkeley National Laboratory, Berkeley, CA, USA; 0000-0002-2939-5398

Nikos C. Kyrpides; US Department of Energy Joint Genome Institute, Lawrence Berkeley National Laboratory, Berkeley, CA, USA; 0000-0002-6131-0462

Janet K. Jansson; Biological Sciences Division, Pacific Northwest National Laboratory, Richland, WA USA; 0000-0002-5487-4315

Jason E. McDermott; Biological Sciences Division, Pacific Northwest National Laboratory, Richland, WA USA; 0000-0003-2961-2572

Kirsten S. Hofmockel; Biological Sciences Division, Pacific Northwest National Laboratory, Richland, WA USA; 0000-0003-1586-2167

National Ecological Observatory Network; NEON, Boulder, CO USA; none; none Jeffrey L Blanchard; Biology, University of Massachusetts Amherst, Leverett, MA USA; 0000-0002-7310-9678

Xiao Jun A. Liu; Institute for Environmental Genomics, University of Oklahoma, Norman, OK, USA; 0000-0002-0488-8591

Jorge L. Mazza Rodrigues; Department of Land, Air and Water Resources, University of California - Davis, Davis, CA, USA; 0000-0002-6446-6462

Zachary B. Freedman; Department of Soil Science, University of Wisconsin-Madison, Madison, WI, USA; 0000-0001-9160-7470

Petr Baldrian; Laboratory of Environmental Microbiology, Institute of Microbiology of the Czech Academy of Sciences, Prague, Czech Republic; 0000-0002-8983-2721

Martina Stursova; Laboratory of Environmental Microbiology, Institute of Microbiology of the Czech Academy of Sciences, Prague, Czech Republic; 0000-0003-1387-6426

Kristen M. DeAngelis; Microbiology Department, University of Massachusetts Amherst, Amherst, MA, USA; 0000-0002-5585-4551

Sungeun Lee; Ecole Centrale de Lyon, Ecully, France; 0000-0001-8878-6463

Filipa Godoy-Vitorino; Department of Microbiology and Medical Zoology, University of Puerto Rico School of Medicine, Medical Sciences Campus, PO Box 365067, 00936 San Juan, Puerto Rico, USA; 0000-0003-1880-0498

Yun Kit Yeoh; Australian Institute of Marine Science, Townsville, Queensland, Australia; 0000-0003-0241-6117

Hinsby Cadillo-Quiroz; School of Life Sciences, Arizona State University; 0000-0002-4908-4597

Susannah G. Tringe; Environmental Genomics and Systems Biology, Lawrence Berkeley National Laboratory, Berkeley, CA, USA; 0000-0001-6479-8427

Archana Chauhan; Panjab University, Chandigarh, India; 0000-0002-6821-0464

Don A Cowan; Centre for Microbial Ecology and Genomics, Department of Biochemistry, Genetics and Microbiology, University of Pretoria, Pretoria, South Africa; 0000-0001-8059-861X

Marc W. Van Goethem; Centre for Microbial Ecology and Genomics, Department of Biochemistry, Genetics and Microbiology, University of Pretoria, Pretoria, South Africa; 0000-0001-8688-3323

Tanja Woyke; U.S. Department of Energy Joint Genome Institute, Lawrence Berkeley National Laboratory, Berkeley, California, USA; 0000-0002-9485-5637

Nicholas C. Dove; University of California, Merced--Merced, CA USA; 0000-0003-1152-956x

Konstantinos T. Konstantinidis; School of Civil and Environmental Engineering, and School of Biological Sciences, Georgia Institute of Technology, Atlanta, GA, USA.; 0000-0002-0954-4755

Thomas E. Juenger; Department of Integrative Biology, University of Texas, Austin, Texas; 0000-0001-9550-9288

Stephen C. Hart; Department of Life and Environmental Sciences and the Sierra Nevada Research Institute, University of California, Merced, CA, USA; 0000-0002-9023-6943

David D. Myrold; Department of Crop and Soil Science, Oregon State University, Corvallis, OR, USA; n/a; n/a

Tullis C. Onstott; Department of Geosciences, Princeton University, Princeton, NJ, USA; n/a; n/a

Brendan J.M. Bohannan; Institute of Ecology and Evolution, University of Oregon, Eugene, OR, USA; 0000-0003-2907-1016

Marty R. Schmer; United States Department of Agriculture, Agricultural Research Service, Lincoln, NE; 0000-0002-3721-6177

Nathan A. Palmer; Wheat, Sorghum, and Forage Research Unit. United States Department of Agriculture - Agricultural Research Service. Lincoln, NE. USA; 0000-0001-8088-6371

Klaus Nusslein; Department of Microbiology, University of Massachusetts - Amherst, MA; 0000-0002-0663-4448

Thulani P. Makhalanyane; DSI/NRF SARChI in Marine Microbiomics, Department of Biochemistry, Genetics and Microbiology, University of Pretoria; 0000-0002-8173-1678

Katherine A. Dynarski; Department of Land, Air, and Water Resources, University of California Davis, Davis, California, USA; 0000-0001-5101-9666

Neslihan Tas; Climate and Ecosystem Science Division, Lawrence Berkeley National Laboratory, Berkeley, CA; 0000-0001-7525-2331

Graeme W. Nicol; Laboratoire Ampère, Ecole Centrale de Lyon, Ecully, France; 0000-0002-3876-022X

Christina Hazard; Laboratoire Ampère, Ecole Centrale de Lyon, Ecully, France; 0000-0002-0325-5856

Erin D. Scully; USDA-ARS Center for Grain and Animal Health Research Manhattan, KS USA; 0000-0002-8315-5619

Kunal R. Jain; Environmental Genomics and Proteomics Lab, Department of Biosciences, Satellite Campus, Vadtal Raod, Sardar Patel University, Bakrol - 388 315 (Anand), Gujarat, India; 0009-0001-6346-9055

Datta Madamwar; P. D. Patel Institute of Applied Sciences, Charotar University of Science and Technology, Changa - 388 421 (Anand), Gujarat, India; 0000-0003-3301-1120

Andrew Bissett; Commonwealth Scientific and Industrial Research Organisation, Hobart, Tasmania, Australia, 7000; 0000-0001-7396-1484

Philippe Constant; Centre Armand-Frappier Santè Biotechnologie, Institut national de la recherche scientifique, Laval, Quèbec, Canada; 0000-0003-2739-2801

Rafael S. Oliveira; Department of Plant Biology, University of Campinas, Campinas, São Paulo, Brazil; 0000-0002-6392-2526

Cristina Takacs-Vesbach; Department of Biology, University of New Mexico, Albuquerque, New Mexico; 0000-0002-5535-2201

Melissa A. Cregger; Biosciences Division, Oak Ridge National Laboratory, Oak Ridge TN; 0000-0001-8329-366X

Alyssa A. Carrell; Biosciences Division, Oak Ridge National Laboratory, Oak Ridge, TN, USA; 0000-0003-1142-4709

Dawn M. Klingeman; Oak Ridge National Laboratory. Oak Ridge TN 37831; 0000-0002-4307-2560

Nicole Pietrasiak; School of Life Sciences, University of Nevada Las Vegas, Las Vegas, NV, USA; 0000-0003-4636-8006

