## Supplementary material for "A global atlas of soil viruses reveals unexplored biodiversity and potential biogeochemical impacts": SI

### SUPPLEMENTAL FIGURES AND TABLES

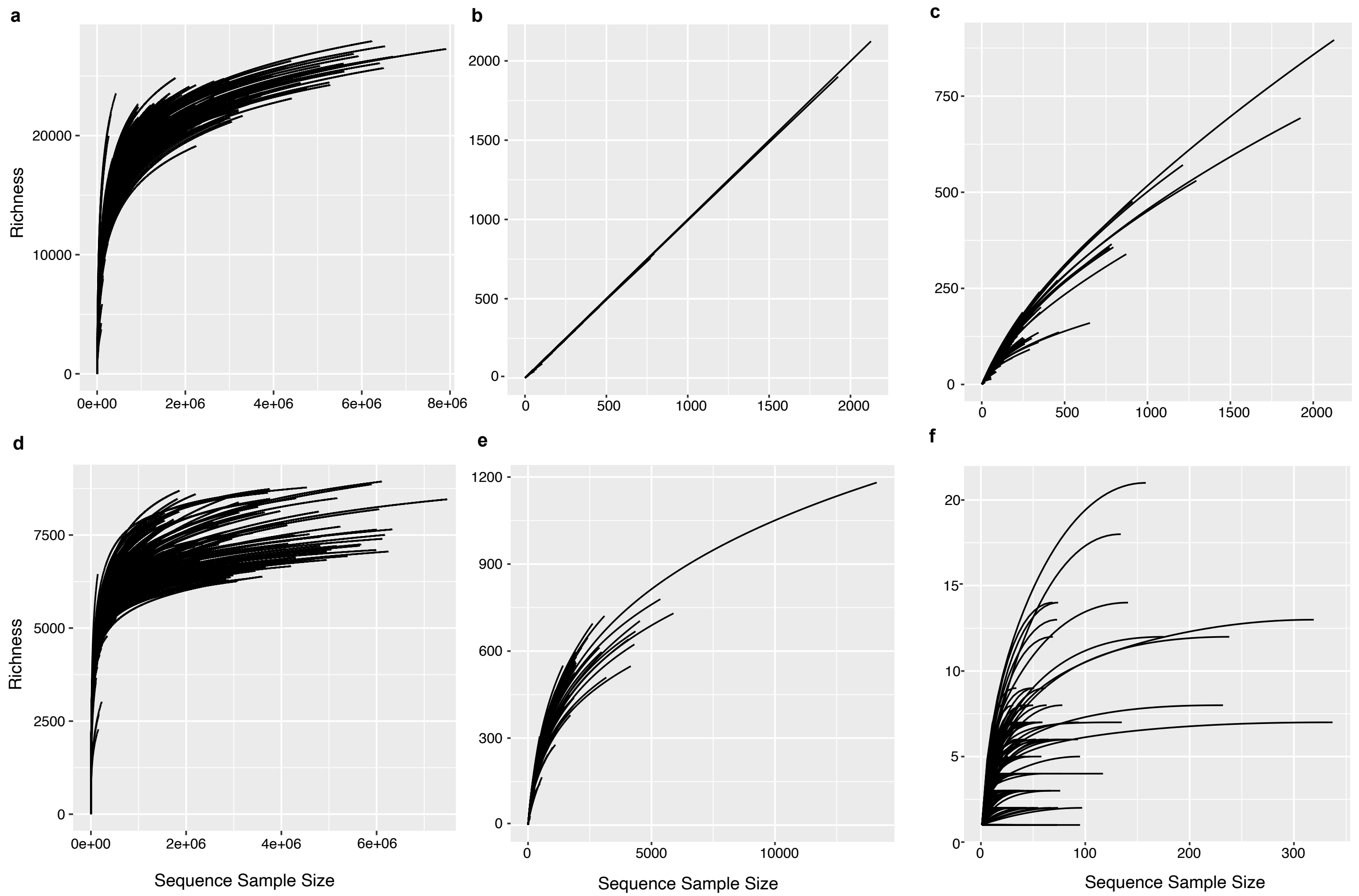


**Figure S1. Rarefaction curves. a,** taxonomy from whole metagenomic sequences, **b**, vOTUs at the species level, **c**, viral clusters at the family level, **d,** Pfams from whole metagenomic sequences, **e**, Pfams from UViGs, **f**, CRISPR spacer-based host assignment of uViGs.

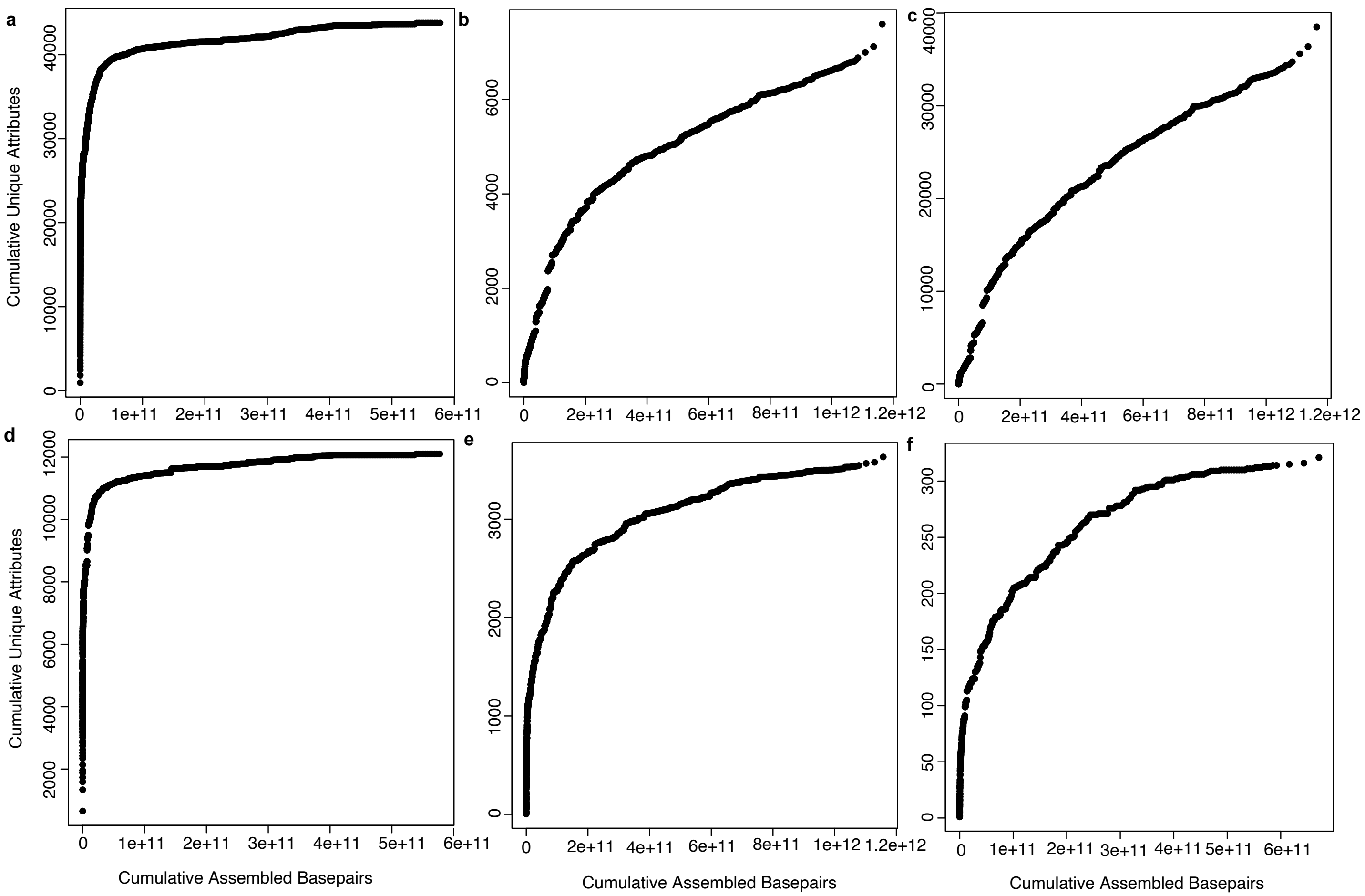


**Figure S2. Total sequencing depth versus cumulative unique attributes. a,** taxonomy from whole metagenomic sequences, **b**, vOTUs at the species-level, **c**, viral clusters at the family-level, **d,** Pfams from whole metagenomic sequences, **e**, Pfams from UViGs, **f**, CRISPR-based host assignment of UViGs.

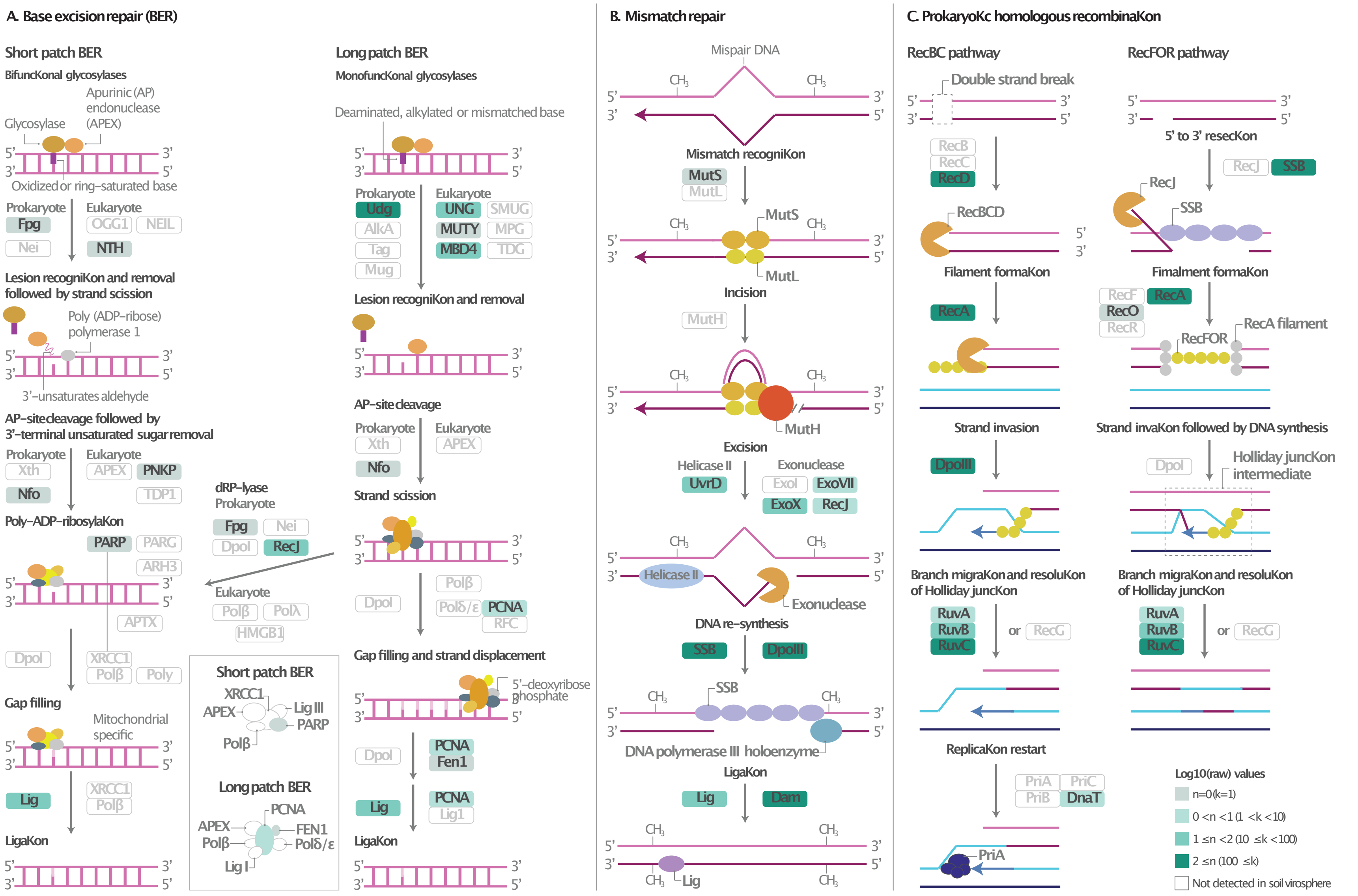


**Figure S3. Metabolic potential encoded by the soil virosphere, hallmarks of viral activity. a,** base excision repair (map03410), **b,**

(prokaryotic) homologous recombination (map03440), and **c**, (prokaryotic) DNA mismatch repair (map03430). KEGG pathways are cropped and/or simplified to enhance visualization. Graphics are adapted from visualizations rendered by Pathview. Color scale denotes the log10 of the total abundance across the entire soil virosphere.

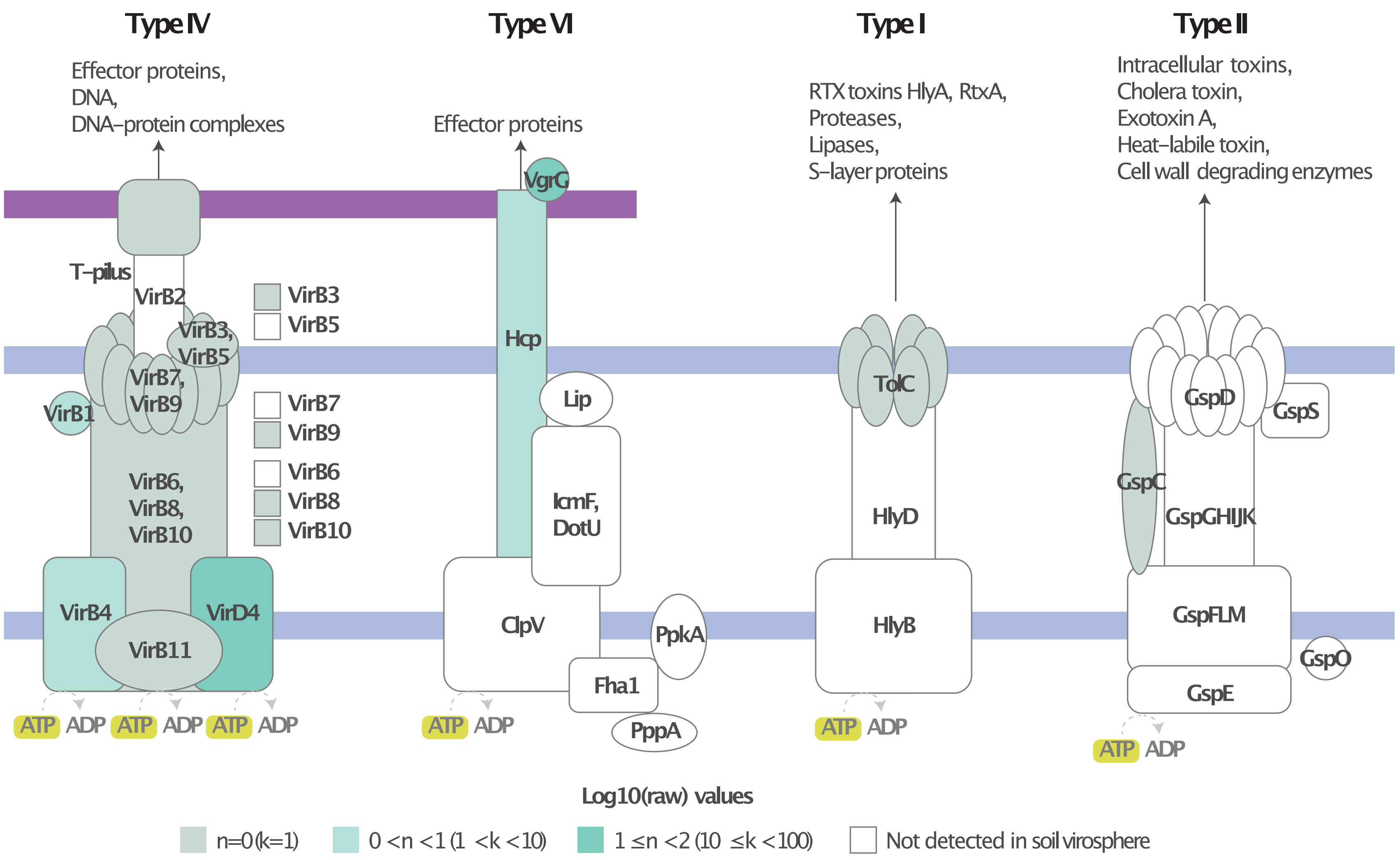


**Figure S4. Metabolic potential encoded by the soil virosphere, bacterial secretion systems (map03070).** KEGG pathways are cropped and/or simplified to enhance visualization. Graphics are adapted from visualizations rendered by Pathview. Color scale denotes the log10 of the total abundance across the entire soil virosphere.

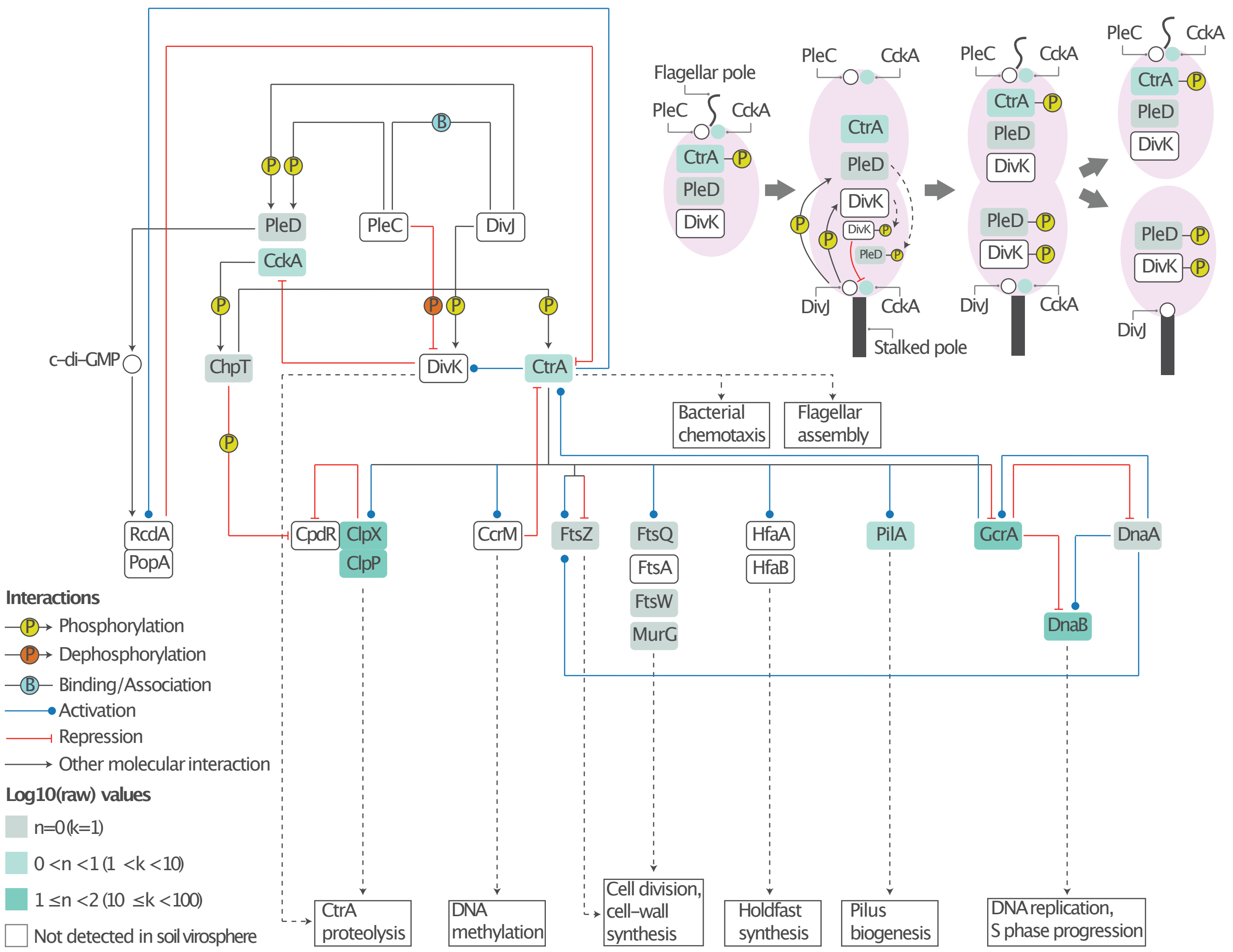


**Figure S5. Metabolic potential encoded by the soil virosphere, *Caulobacter* cell** cycle (map04112). KEGG pathways are cropped and/or simplified to enhance visualization. Graphics are adapted from visualizations rendered by Pathview. Color scale denotes the log10 of the total abundance across the entire soil virosphere.

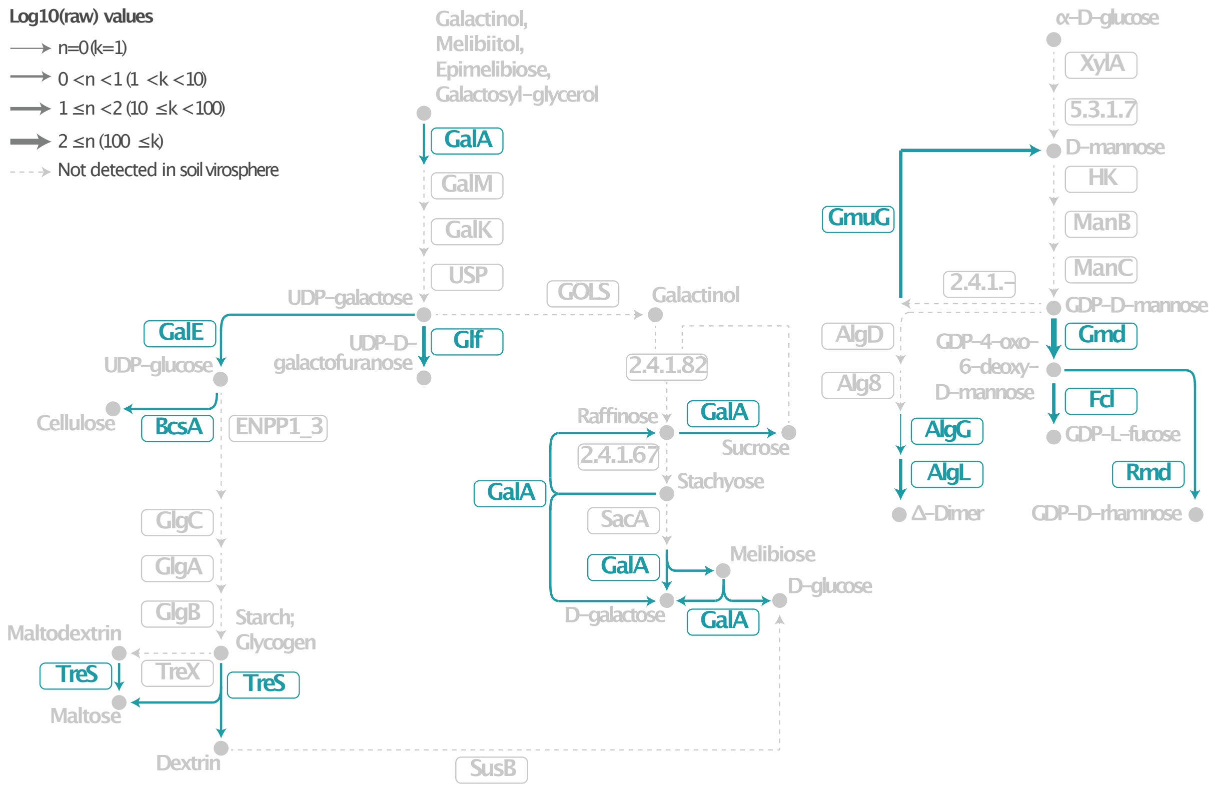


**Figure S6. Metabolic potential encoded by the soil virosphere, hallmarks of viral activity.** Portions of galactose metabolism (map00052), starch and sucrose metabolism (map00500), and fructose and mannose metabolism (map00051) are depicted. KEGG pathways are cropped and/or simplified to enhance visualization. Graphics are adapted from visualizations rendered by Pathview. Color scale denotes the log10 of the total abundance across the entire soil virosphere.
